# Stoichiometry-dependent specificity in biotin enrichment: a benchmarking framework for proximity labeling proteomics

**DOI:** 10.64898/2026.05.07.723439

**Authors:** Christopher Árpád Zala, Maria Cristina Trueba Sanchez, Jelle van den Bor, Tijmen Willemsens, Frederik Johannes Verweij, Maarten Altelaar, Kelly Stecker

## Abstract

Proximity labeling methods (including, BioID, TurboID, ultraID), along with surface proteomics and microdomain mapping, enable proteome-wide identification of spatially proximal proteins via MS-based analysis. These workflows require specific enrichment of biotinylated proteins using affinity purification, yet enrichment specificity can often be compromised by non-specifically bound proteins. As labeling strategies are increasingly applied to complex biological samples with low protein input or low biotin stoichiometry, accurately distinguishing true targets from background becomes a major analytical challenge. Despite its critical impact on data quality and interpretation, the influence of biotinylation level and protein input on enrichment performance remains poorly characterized, limiting the reliability of proximity labeling experiments.

To address this, we establish a quantitative benchmarking framework that systematically evaluates biotin enrichment under controlled conditions, including scenarios of low biotin stoichiometry. Using this setup, we show that enrichment specificity strongly depends on biotin stoichiometry: higher levels of biotinylation in samples yield high specificity, whereas low biotinylation increases non-specific background. Reduced protein input further limits recovery of true targets, yet maintains enrichment specificity, highlighting sensitivity constraints of enrichment-based workflows.

We apply this framework to biotinylated extracellular vesicle (EV) cargo uptake in recipient cells using ultraID-CD63 labeling. Detection of the most abundant EV cargo proteins under low biotinylation conditions indicates that current workflows approach the lower bounds of biotin enrichment sensitivity. Together, these standards provide a practical reference for evaluating and optimizing biotin enrichment workflows, supporting quantitative and reproducible proximity labeling in proteomics.

## Introduction

Modern proteomics aims to comprehensively characterize protein function, localization and interactions within cells^1^. Understanding how proteins localize and interact inside cells and on the cell surface is essential for deciphering cellular processes and disease mechanisms. Identifying interacting subsets of proteins remains a challenging task, especially when those proteins are localized to specific subcellular niches that are difficult to isolate without disrupting their native context ^2^. To address these limitations, biotin-based labeling approaches have been developed to preserve information about subcellular context while enabling specific isolation of target proteins ^3^. Biotin labeling in combination with biotin enrichment and MS analysis provides a powerful workflow to identify subsets of proteins in defined cellular contexts. This strategy underlies several widely used experimental approaches, including: (i) proximity labeling methods (e.g., BioID, APEX, TurboID, ultraID [uID]) for identifying proteins in close spatial proximity; (ii) surface biotinylation techniques for mapping cell surface proteomes across diverse cell types and clinically relevant samples; and (iii) microdomain mapping approaches for characterizing the dynamic landscape of surface receptors and their native protein environment^3–9^. Together, these biotin-based approaches have greatly expanded the ability of proteomics to probe the spatial organization and dynamics of the proteome.

Biotin enrichments are advantageous for several reasons. Biotin binds tightly to streptavidin and other biotin binding proteins such as neutravidin and avidin, allowing for the efficient capture of biotinylated proteins^10^. When immobilized on beads or similar matrices, these reagents allow convenient isolation of biotinylated proteins from complex biological samples^11^. Leveraging the strength of the streptavidin-biotin interaction, researchers can combine biotin enrichment at either protein or peptide level with MS-based proteomics to retrieve biotinylated target proteins from complex samples^12^. To maximize sensitivity and avoid the challenge of efficiently eluting biotinylated peptides, biotin enrichments are most often performed at the protein level^13^. Although this reduces protein input requirements, it presents several technical challenges in identifying true target proteins. First, there is no detection, and thus direct evidence, of biotin tags. Second, biotin enrichments performed at the protein level are more prone to non-specific binding to the enrichment matrix, as the larger size of proteins increases the likelihood of unspecific interactions^13^. For these reasons, downstream data analysis and hit validation in biotin enrichment experiments can be challenging, as the confidence that any detected proteins is a true target is often uncertain.

With the expansion of these approaches, biotin-based enrichment workflows are now being extended to increasingly complex and biologically relevant applications such as intercellular labeling of transiently interacting cells or labeling protein interactions *in vivo*. These settings place substantially greater analytical demands on enrichment performance, as they are often characterized by limited protein input and low biotin stoichiometries. Under such conditions, the fraction of biotinylated target proteins relative to background can become extremely small, making their specific isolation and reliable detection particularly challenging. In this context, it is critical to understand how well biotin enrichment workflows perform near their sensitivity limits and to what extent observed signals reflect true biotinylation rather than background binding. Achieving predictable and interpretable outcomes therefore requires a systematic understanding of how enrichment performance is influenced by biotinylation levels and protein input.

Here, we systematically assess how biotin stoichiometry and protein input influence enrichment quality using defined biotinylated protein standards. We establish a quantitative benchmarking framework that enables controlled evaluation of enrichment performance using a mixed species proteome setup. We demonstrate that biotin stoichiometry is the primary determinant of enrichment performance and provide quantitative insights that form a basis for optimizing biotin-based proteomics workflows, enabling more informed experimental design and data interpretation applicable to diverse biotin labelling applications. To evaluate the biological relevance of our framework, we apply it to a compelling and analytically demanding test case: the proteomic detection of EV cargo transfer between cells. EVs are key mediators of intercellular communication, yet direct biochemical evidence of which proteins are functionally delivered to recipient cells remains elusive, making EV cargo transfer strong test of enrichment performance under biologically limiting conditions^14^.

## Results

### A benchmarking framework for evaluating biotin enrichment quality

In workflows that combine biotin labeling, biotin enrichment, and subsequent MS analysis, the overall quality of the experiment depends on the specificity of the enrichment step. In biotin-based labeling experiments, however, it is not possible to reliably determine whether the number of retrieved proteins reflects the extent of biotin labeling or the performance of biotin enrichment. To systematically assess how biotin stoichiometry and the amount of protein input used affect the specificity of biotin enrichments, we designed a quantitative benchmarking framework to systematically evaluate biotin enrichment performance under controlled conditions. To this end, we chemically biotinylated *E. coli* protein lysates and mixed them at different percentages into human background protein lysates (Figure 1A), generating five biotinylated protein standards with differing biotin levels. This approach offers several advantages: (i) genomic differences between human and *E. coli* proteins enable clear distinction between target and background proteins by MS, circumventing the need for direct biotinylation detection; (ii) the experimental setup allows precise control of target abundance and total protein input, enabling systematic assessment of the effect of biotin stoichiometry and protein input on enrichment performance; and (iii) the use of a biotinylated *E. coli* proteome within a human proteome background approximates the labeling complexity and non-specific binding observed in biotin-based labeling experiments in human cells. Together, these features provide a controlled yet biologically relevant system for assessing the factors that influence biotin enrichment performance.

**Figure 1.**
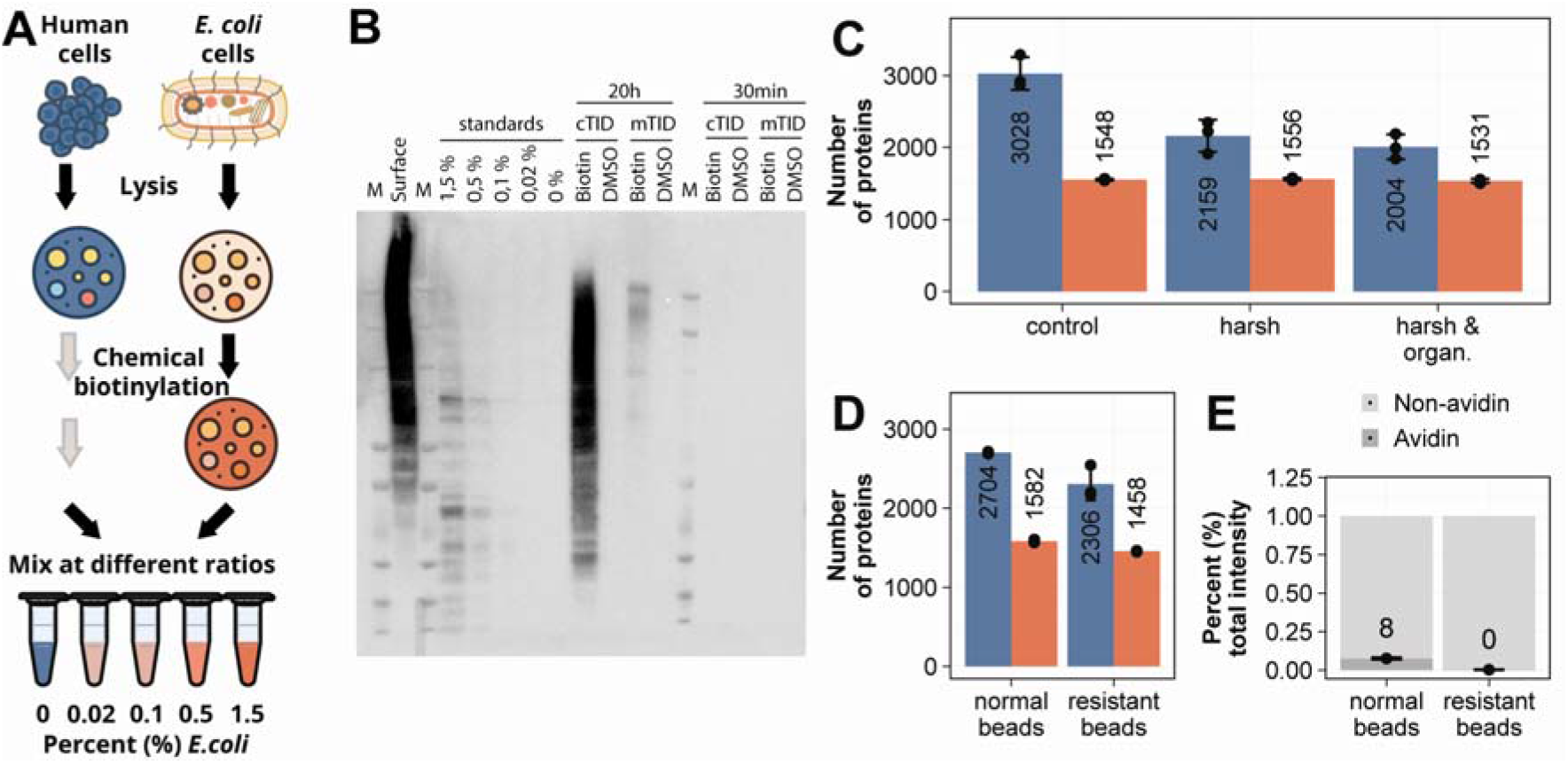
Biotinylated protein standards reflect labeling regimes encountered in practice and optimized enrichment workflow decreases non-specific background binding. **A)** Schematic illustrating the experimental system. *E. coli* cells are lysed and their proteins chemically biotinylated. The biotinylated *E. coli* lysate is then mixed at defined ratios with an unmodified human protein lysate to generate samples with varying target-to-background compositions. **B)** Western blot of cell lysates of surface-labeled cells and of cytosolic (cTID) or membrane-localized TurboID (mTID) cell lysates after 20 h or 30 min labeling, alongside biotinylated benchmark standards. DMSO lanes indicate negative control samples treated with dimethyl sulfoxide (DMSO) but not biotin. Each lane contains 20 µg of the respective sample. Biotinylated proteins were detected with streptavidin–HRP. **C)** Bar plot showing the number of *E. coli* proteins (red, target) or human proteins (blue, background) detected by LC-MS after enrichment of 0.1% biotinylated *E. coli* protein lysate in 500ug of human background with Pierce agarose beads across different wash conditions. Control wash contains low percentages of detergent, harsh wash contains high percentages of detergent, harsh & organic wash contains high percentages of detergent with subsequent organic solvent wash. **D)** Bar plot showing the number of *E. coli* (red, target) or human (blue, background) proteins detected by LC-MS after enrichment of 0.5% biotinylated *E. coli* lysate in 250ug of human background with either normal Pierce agarose beads or in-house made protease resistant beads. **E)** Stacked bar plot showing the percentage of total MS intensity attributed to avidin (dark grey) and all remaining proteins (light grey) following enrichment using either standard Pierce agarose beads or in-house–produced protease-resistant Pierce agarose beads.

To calibrate this benchmarking framework to biotin labeling levels encountered in conventional samples, we compared the biotin content of our standards to those observed in commonly used biological labeling approaches. We analyzed samples with high biotin stoichiometry: surface glycan labeling, generated by alkoxyamine-PEG4-mediated biotin labeling, and cytosolically expressed TurboID with extended biotin incubation (20h)^4,7^. For low biotin stoichiometry we tested cytosolic TurboID with short biotin incubation (30 min) and membrane-localized TurboID with both 20h and 30min biotin incubations. Cytosolic TurboID localizes to the cytoplasm and labels intracellular proteins broadly, whereas membrane-localized TurboID, a GPI-anchored variant, predominantly labels plasma membrane and secretory pathway proteins. Western blot analysis (Figure 1B) revealed that surface glycan labeling and prolonged (20 h) cytosolic TurboID exceeded the highest standard (∼1.5% biotinylation), whereas both membrane-localized TurboID and cytosolic TurboID with 30min of labeling yielded lower labeling levels (0.1-0.5% biotinylation) (Figure 1B, Supplementary Figure 1A). Finally, membrane-localized TurboID with 30 min of labeling matched the lowest levels of biotinylation standards of ∼0.02% biotin. These comparisons position our standards within the dynamic range of biologically relevant experiments and provide a quantitative reference for estimating labeling efficiency and required protein input.

### Optimizing the enrichment workflow boosts biotin enrichment quality

Having established both a controlled benchmarking system, we next leveraged this framework to optimize the enrichment workflow itself by evaluating several parameters: washing conditions, resin selection, and resin derivatization to resist protease digestion.

Insufficient stringency during bead-based enrichment can lead to substantial non-specific protein binding. To make sure that we minimize non-specific binding through insufficient wash conditions, we tested whether increasing the stringency of the wash conditions routinely used in our laboratory could improve the specificity of the biotin enrichment. We performed a biotin enrichment of the biotinylated standard sample at 0.1% biotinylation in a total of 500 µg of protein input. We compared the wash conditions routinely used in the laboratory consisting of low detergent washes of the beads (0.1% SDS), with one previously published method where high detergent buffers were applied (2% SDS)^15^. Additionally, we supplemented the high detergent wash protocol with extra organic solvent washes (30% ACN) as a third condition. When evaluating the number of background proteins present in each enrichment, we saw that high percentages of detergent drastically decreased the number of background proteins relative to low percentage of detergent (Figure 1C). Adding organic solvent washes to the high detergent washes further decreased the number of background proteins. The number of target proteins stayed stable over all wash conditions. We found that high detergent washes in combination with organic solvent washes lead to the strongest reduction of background proteins and therefore we used these wash conditions in all following experiments.

There are many different suppliers offering bead-based biotin enrichment matrices coupled to biotin binding proteins. Since it is clear in the field that different bead types have different biotin binding capacities, we anticipate that this parameter influences the outcome of biotin enrichments ^16^. We tested the biotin binding capacity of 3 different bead types (Pierce agarose neutravidin, Pierce magnetic streptavidin, Resyn magnetic streptavidin) on our biotinylated standard sample. We used 500 µg total input sample with the highest biotinylation level (1.5% biotinylated *E. coli* proteins) and tested different bead amounts. Western blot analysis revealed that the three bead types depleted biotin from the sample at different bead volumes (Supplementary Figure 1B). While for the Pierce agarose beads, 5 µl of beads was sufficient to deplete all biotin from the samples, for Pierce magnetic beads and Resyn beads 20 µl of beads were required. Since the biotin depletion efficiency of the three bead types differed, we evaluated the effect on the efficiency of biotin enrichments. We enriched the 1.5% biotinylated standard sample in a total input amount of 500 µg protein and analyzed the samples by MS. We used 10 µl of beads for agarose and 20 µl for Pierce and Resyn, respectively. For pipetting consistency, we had to increase the bead volume of agarose beads from 5 µl to 10 µl. The recovered number of target proteins varied across bead types. The most target (*E. coli*) proteins were recovered using pierce agarose beads (1348 proteins), followed by Pierce magnetic beads (849) and Resyn beads (750). Surprisingly, Resyn beads also yielded a much larger number of background proteins, leading us to exclude them from subsequent evaluations (Supplementary Figure 1C).

To release proteins from the beads after biotin enrichment, on-bead digestion is commonly performed. However, this process also digests the biotin-binding protein immobilized on the bead matrix (e.g., streptavidin, neutravidin). Because these biotin-binding proteins are present in large quantities, digestion generates highly abundant bead-derived peptides that dominate MS spectra and reduce the dynamic range available for detection of true analytes. Previously, a method was introduced to make streptavidin on beads resistant against proteolytic cleavage^17^. We tested whether making beads resistant against protease digestion and thereby reducing overloading in the mass spectrometer due to bead-derived peptides could improve target protein recovery. We enriched our 0.1% biotinylated standard sample in a total protein input of 500 µg with either unmodified agarose beads or in-house modified protease resistant agarose beads. Even though the protease resistant beads reduced contamination by bead-derived peptides efficiently, we did not see an increase in the number of target proteins (Figure 1 D&E). On the contrary, the number of retrieved target proteins was slightly decreased. Due to this slight decrease in recovered target proteins, we decided to use unmodified beads for future experiments.

### Biotin enrichment performance is dependent on biotinylation level in the samples

Next, we wanted to assess how biotin stoichiometry influences enrichment efficiency in terms of two key parameters: percentage of total protein abundance of target proteins (*E. coli*) and the number of retrieved target proteins. We performed this assessment in comparison between two of the previously tested bead types, Pierce agarose and Pierce magnetic beads. In the comparison, we kept the input protein amount constant at 500 µg and included the full range of our biotinylated standard samples (0% - 1.5% biotin). We then enriched the samples separately with the two bead types and subjected them to MS analysis. We observed that when enriching with agarose beads, we could allocate a higher percentage of the total protein abundance in the sample to target proteins relative to magnetic beads across all biotinylation percentages of the samples (Figure 2A&B). Furthermore, agarose beads yielded higher numbers of target proteins and lower number of background proteins across all biotinylation percentages of samples compared to magnetic beads (Figure 2C&D). These findings indicate that agarose beads outperformed magnetic beads. For both bead types we found that the number of target proteins systematically decreased with decreasing biotin stoichiometry (Figure 2C&D). The percentage of the total protein abundance of target proteins is similarly decreasing (Figure 2A&B). Both bead types efficiently enriched biotinylated *E. coli* proteins relative to unenriched controls, confirming effective enrichment performance. (Figure 2E&F). Due to the superior performance of agarose beads compared to magnetic beads, we decided to focus on agarose beads in the following experiments and analysis.

**Figure 2.**
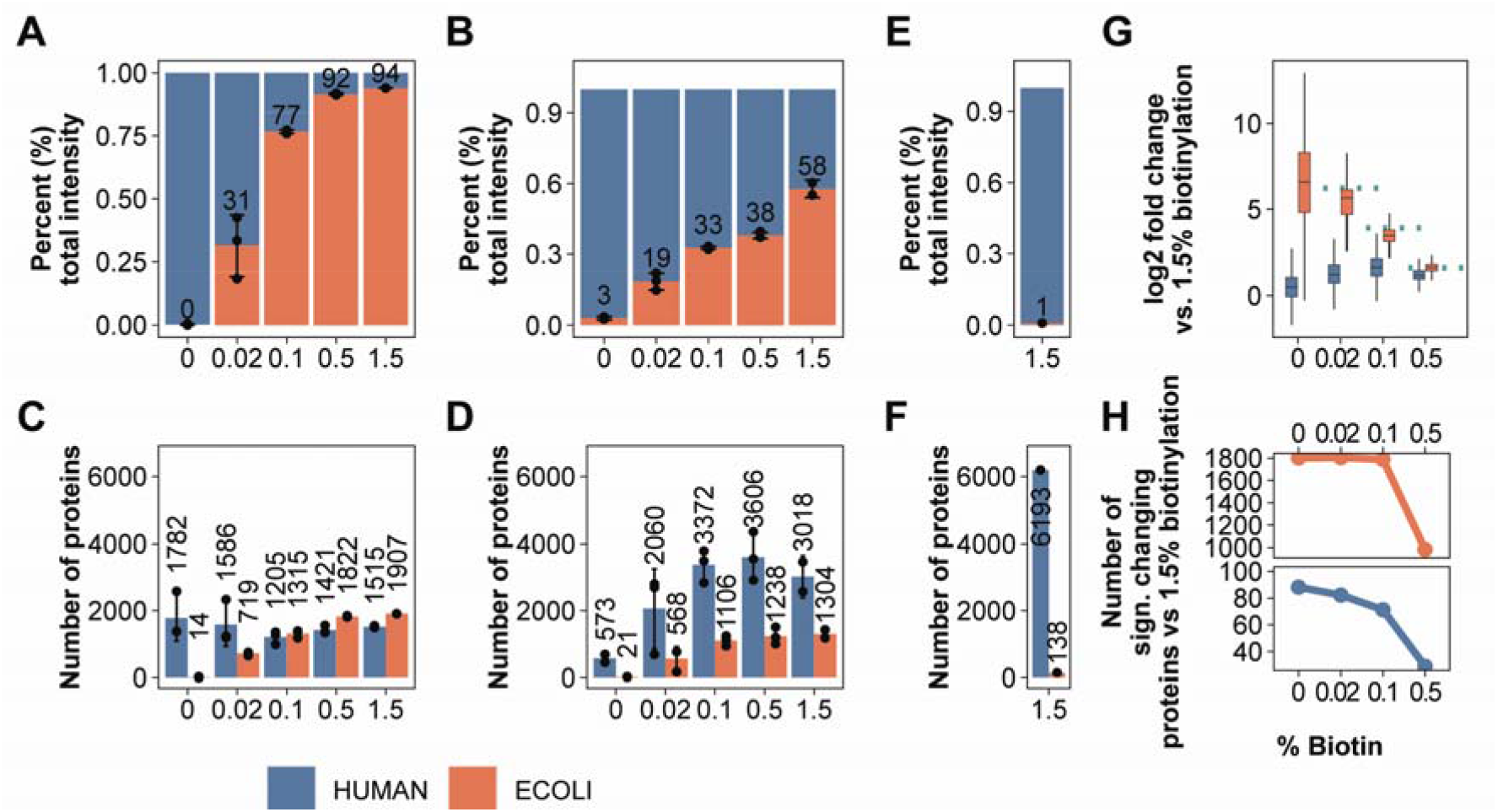
Agarose beads outperform pierce magnetic beads in enrichment specificity and in number of retrieved target proteins. **A, B, E)** Stacked bar plots showing the percentage of total MS intensity attributed to *E. coli* (red, target) and human (blue, background) proteins at different biotinylation levels. Panels compare enrichment using (A) Pierce agarose beads, (B) Pierce magnetic beads, and (E) no enrichment. **C, D, F)** Bar plots showing the number of E. coli (red, target) or human (blue, background) proteins identified at different biotinylation percetages. Panels compare enrichment using (C) Pierce agarose beads, (D) Pierce magnetic beads, and (F) no enrichment. **G)** Box plots showing fold change of *E. coli* (red, target) and human (blue, background) proteins. Biotinylation percentages are compared to the 1.5% biotinylated sample. The green dotted line indicates the theoretical fold change based on the ratio of biotinylation content between samples. **H)** Line plots showing the number of significantly changing E. coli (red, target) and human (blue, background) proteins across biotinylation percentages (x-axis), relative to a 1.5% biotinylation reference. Significance was determined using a fold-change threshold of 2, except at 0.5% biotinylation (asterisk), where a threshold of 1 was applied. Y-axis is broken to show protein numbers of human and *E. coli* proteins more accurately. Number of significantly changing proteins are derived from differential abundance analysis displayed in Supplementary Figure 2A.

Next, we assessed the quantitative robustness of the biotin enrichment workflow. Because samples were generated with defined biotinylation levels, the relative abundance of enriched proteins was expected to reflect the theoretical ratios between these conditions. To test this, we compared samples enriched with agarose beads at 1.5% biotinylation to all other biotinylation levels using pairwise statistical analysis. As anticipated, most *E. coli* proteins displayed higher abundance in the 1.5% condition (Supplementary Fig. 2A).

Consistent with this observation, the measured enrichment ratios closely matched the theoretical ratios expected from the corresponding biotinylation levels (Figure 2G). In contrast, the number of significantly changing human background proteins remained largely constant across all comparisons, indicating that the enrichment of false positive human proteins is largely independent of sample biotinylation levels (Figure 2H). This trend is corroborated by a consistently stable false discovery rate of 3–5%. These results show that quantitative changes in biotinylated target proteins are accurately captured by the enrichment workflow, whereas false-positive identifications appear to remain largely stable across conditions.

### Enrichment efficiency remains stable across reduced input protein amounts

In the experiments described above, enrichments were performed using a relatively high protein input of 500 µg. Since high protein inputs are not always available in biological experiments, we investigated how decreasing the protein input amount affects the enrichment efficiency. We used the full range of our biotinylated protein standards (0 – 1.5%) while we decreased protein input amounts to 300, 100 and 50 µg. We again assessed the percentage of total protein abundance of target proteins and the number of retrieved target proteins (Figure 3A&B). As expected, the number of retrieved target proteins decreased with decreasing input amount (Figure 3B). Surprisingly, however, the percentage of total protein abundance of target stayed stable over different input amounts (Figure 3A). These results indicate that while the number of target analytes decreases with decreasing protein input amount, the efficiency of the enrichment in terms of percentage of retrieved target proteins is stable over different inputs.

**Figure 3.**
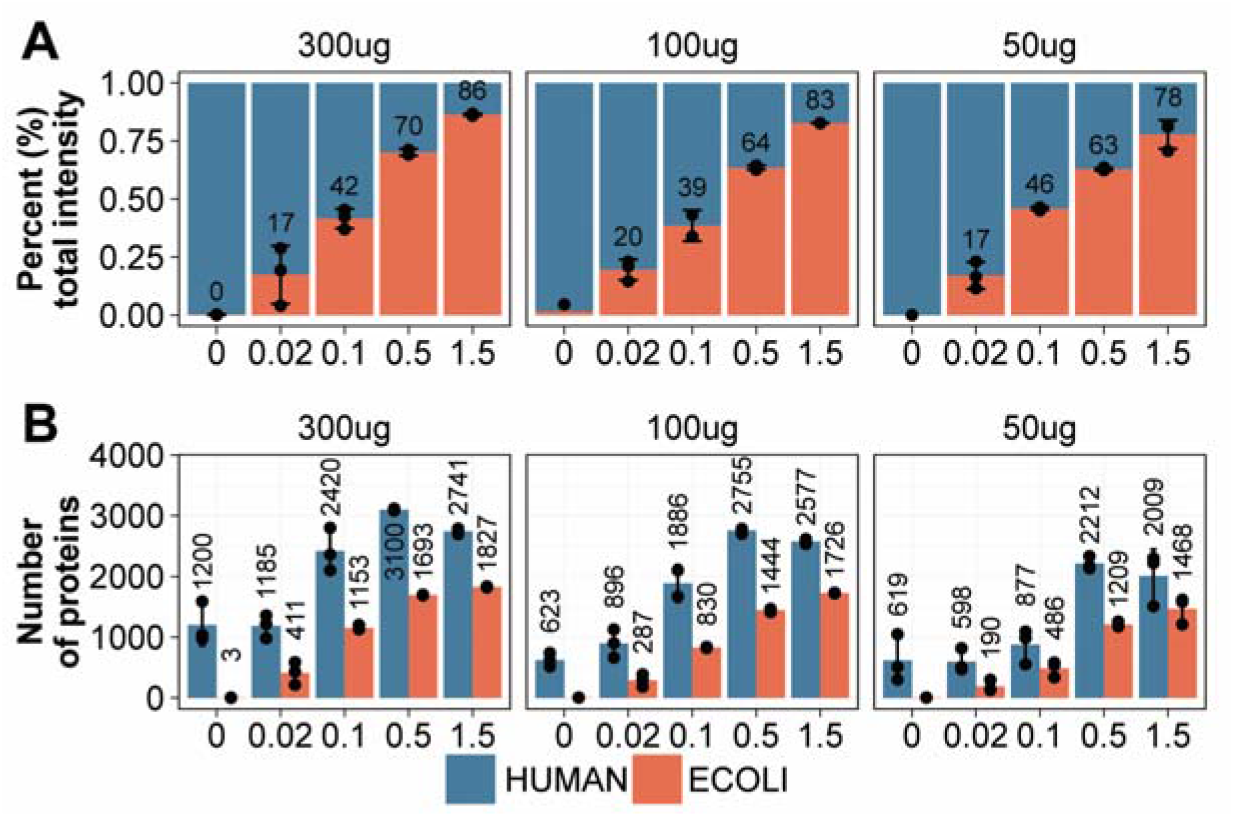
Decrease of total protein input amount leads to loss of target proteins but does not affect the enrichment efficiency. **A)** Stacked bar plot showing the percentage of total MS intensity attributed to *E. coli* (red, target) and human (blue, background) proteins at different biotinylation percentages across varying protein inputs. **B)** Bar plot showing the number of *E. coli* (red, target) and human (blue, background) proteins detected by LC-MS at different biotinylation percentages across varying protein inputs.

### Biotin enrichment standards are reflected in EV derived samples with different biotinylation levels

Whether this relationship between protein input, labeling extent, and enrichment performance is maintained in complex biological systems remains an important question. EVs are key mediators of intercellular communication. However, identifying proteins that are functionally transferred to recipient cells is inherently challenging. EV-derived proteins must be detected against an abundant endogenous proteome, making it difficult to distinguish transferred cargo from background without selective labeling strategies. This challenge is further compounded by the typically low efficiency of EV uptake, which limits the detectable signal^18^. As a result, the identity of proteins that are functionally delivered and retained remains poorly defined, hindering a mechanistic understanding of how EVs influence recipient cell behavior. To address this question, we designed a proximity labeling experiment to trace EV cargo delivered to receiver cells (Figure 4A). HeLa Flp-In cells were genetically engineered with a uID-CD63 construct, placing uID on the luminal side of the canonical EV marker CD63 to enable efficient biotinylation of intraluminal EV proteins. Biotinylated EVs were then incubated with receiver cells, and the retained EV content was analyzed by biotin enrichment and MS. Based on the expected dilution of EV cargo in the recipient cells, we hypothesized that the total biotin content in receiver cell lysates would be low, representing a stringent test of enrichment sensitivity and specificity under challenging biological conditions.

**Figure 4.**
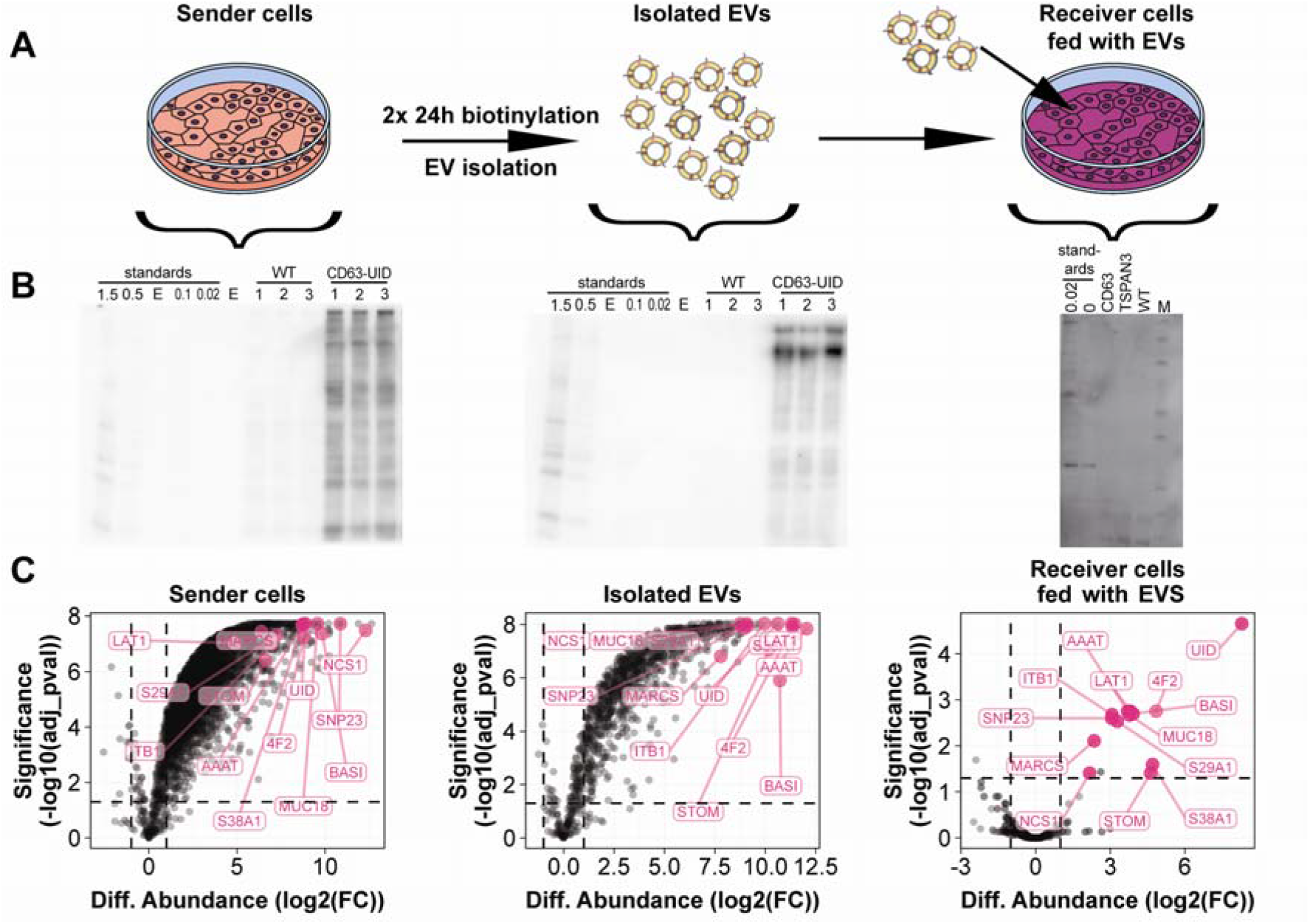
Biotin level of EV samples strongly influences the quality of the enrichment. **A)** Schematic illustrating the experimental system. Biotinylation of EV sender cells is induced by supplying biotin to the medium for 2x 24h. After that EVs are isolated. Isolated EVs are then fed to Hep2G receiver cells to retrieve internalized EV proteins by biotin enrichment. **B)** Westernblot of lystates from respective samples from panel A alongside biotinylated benchmark standards. Each lane contains 20 µg of the respective sample. Biotinylated proteins were detected with streptavidin–HRP. Letter E marks empty lanes. **C)** Volcano plots showing differential protein abundance versus significance (–log10(adjusted p-value)) of respective samples from B after biotin enrichment. The comparison is always between CD63-UID versus WT. Proteins significantly upregulated in receiver cell enrichment are highlighted in pink.

To generate biotinylated EVs, HeLa Flp-In Myc–uID–CD63 cells were supplemented with biotin (Figure 4A), while WT cells served as controls. EVs were isolated by size exclusion chromatography after 24 h of secretion. Both uID–CD63 and WT EVs displayed comparable size distributions, as assessed by dynamic light scattering (Supplementary Figure 3A). EV sender cells were collected separately to be subjected to biotin enrichment and MS analysis. Biotin content of EVs and sender cells was analyzed by western blot in direct comparison to biotinylated protein standards (Figure 4B). Both biotinylated sender cells and biotinylated EVs showed biotin content considerably above the 1.5% biotinylated protein standard (Figure 4B). To identify biotinylated EV proteins, we performed biotin enrichments on lysates of EV sender cells and isolated EVs and analyzed those samples by MS. We saw that in comparison to WT samples, biotinylated sender cells and biotinylated EVs showed increased protein intensities for most proteins after biotin enrichment (Figure 4C). In addition, both biotin-enriched sender cells and EV samples showed strong enrichment of the Gene Ontology term “extracellular exosome,” consistent with their EV origin (Supplementary Figure 4B&C). These findings are in line with the high biotin content observed on WB of biotinylated sender cells and EVs, indicative of a good enrichment quality. Indeed, we observed significant upregulation of biotinylated sender cell protein content and biotinylated EV protein content compared to the respective WT samples by MS analysis after biotin enrichment. We defined true biotinylated EV proteins as proteins passing this significance and differential abundance threshold in comparison to WT EVs. Because the uID-CD63 construct traffics throughout the secretion machinery, sender cells were anticipated to contain more biotinylated proteins than EVs. As expected, EV proteins constituted a subfraction of the biotinylated proteins found in sender cells. (Supplementary Figure 3D). Furthermore, we see both uID and CD63 being among the proteins with highest protein intensities and greatest differential abundance in biotionylated EVs compared to WT EVs after biotin enrichment. Upregulation to a similar degree of uID and CD63 was observed in sender cells after biotin enrichment.

We also analyzed the whole EV lysates by MS. Because merely proteins in close proximity to CD63 are biotinylated in uID-CD63 EVs, the biotinylated proteins detected in EVs likely represent only a subfraction of total proteins present in the EV lysates. (Supplementary Figure 3D). Since the proteomic profiles of biotinylated EVs closely resembled those of WT EVs, the genetic modification of sender cells appears to have minimal effect on EV protein composition. (Supplementary Figure 3E&F). Taken together these findings show that genetically engineered HeLa uID-CD63 cells produce highly biotinylated EVs and subsequent biotin enrichment specifically enriches for biotinylated EV proteins.

After characterizing biotinylated EV content, we set out to apply those EVs in an EV feeding experiment (Figure 4A). With these experiments we aimed to characterize EV proteins absorbed and retained by receiver cells. For this purpose, Hep2G cells were incubated with uID-CD63 containing biotinylated EVs or WT EVs for 24h, after which cells were washed thoroughly and lysed. Biotin content of EV treated receiver cells was analyzed by WB in comparison with the biotin protein standards (Figure 4B). On WB we did not see biotin signal of receiver cell lysates treated with biotinylated EVs, suggesting that the biotin content of the samples was lower than the protein content of the 0.02% protein standard sample, the lowest point of our standard. The starting material for receiver cell biotin enrichment was limited (45 µg), suggesting we would approach the sensitivity limit of the method. Nonetheless, we enriched 45 µg of lysates from receiver cells incubated with either biotinylated uID-CD63 EVs or WT EVs and analyzed the samples by MS. We find a similar number of proteins in receiver cells fed with either uID-CD63 or WT EVs (Supplementary Figure 4A). When comparing protein intensities of receiver cells fed with biotinylated EVs to receiver cells fed with WT EVs we found 14 proteins to be significantly upregulated after biotin enrichment. Out of those 14 proteins, 13 proteins are part of biotinylated EV proteins. The low number of significantly upregulated proteins after biotin enrichment suggested that the method operates near its detection limit. We therefore examined the intensities of the 13 significantly upregulated proteins in EV sender cells and in biotinylated EVs after biotin enrichment and whole-EV proteome analysis. The 13 proteins significantly enriched in receiver cells after EV feeding and biotin enrichment corresponded mostly to the most abundant proteins biotin-enriched sender cells and biotinylated EVs, consistent with the method operating near its detection limit (Figure 4C and Supplementary Figure 4B&C). While many of these proteins were also among the most abundant proteins in whole EV lysates, a few displayed lower intensity ranks (350–958), indicating variability in EV protein representation in enrichment and whole EV lysate analysis (Supplementary Figure 4D). This difference is also reflected by the low correlation of protein intensities in whole EV lysates and biotin enriched uID-CD63 EVs (Supplementary Figure 4E). To assess the impact of further reduced biotinylation on target recovery, we repeated the EV feeding experiment using uID–TSPAN3 biotinylated EVs, which exhibited lower biotinylation levels than uID–CD63 EVs (Supplementary Fig. 5A).

Under these conditions, biotin enrichment of recipient Hep2G cells did not yield any significantly enriched proteins compared to WT EV controls (Supplementary Fig. 5B). Notably, eight of the 13 proteins previously identified as significantly enriched in uID–CD63 EV–treated cells were still detected following uID–TSPAN3 EV treatment. Although present, those proteins fail to reach statistical significance, likely due to increased variability. These results indicate that at very low biotinylation stoichiometry, variability in enrichment efficiency limits the reliable detection of true target proteins.

Among the 14 proteins incorporated and retained in receiver cells following feeding with uID–CD63 biotinylated EVs, uID was identified, but CD63 was absent from the MS dataset. Furthermore, the identified proteins from a biologically coherent group enriched for plasma membrane transport, adhesion, and membrane trafficking functionalities (Supplementary Figure 6). Several hits belong to nutrient transport systems, including SLC7A5 (LAT1), SLC3A2 (CD98/4F2), SLC38A1, SLC29A1, and SLC1A5, suggesting active EV-modulation of amino-acid and nucleoside uptake typical of metabolically active or proliferating cells ^19^. Notably, LAT1 forms a heterodimer with CD98 (SLC3A2), supporting the functional consistency of the dataset ^20^. Additional proteins such as ITGB1, MCAM, and BSG are involved in cell adhesion and surface signaling, which can couple EV to cell surface interactions and metabolic regulation ^21^. Finally, proteins including SNAP23, NCS1, MARCKS, and STOM participate in membrane organization, vesicle trafficking and EV secretion, processes that can possibly regulate transporter localization to EVs ^22–24^. Together, the hitlist is consistent with enrichment of proteins involved in cell-surface transport and trafficking networks supporting EV-mediated modulation of cellular metabolism and signaling. Applying our framework to EV-mediated protein transfer, we found that enrichment performance closely reflected biotinylation stoichiometry in biological samples. High biotin content in sender cells and EVs enabled robust target recovery, whereas low biotin levels in recipient cells limited detection. Accordingly, uID–CD63 EV– treated cells yielded significantly enriched proteins, while the lower biotinylation of uID–TSPAN3 EVs resulted in no detectable differences from WT controls. Despite these limitations, the proteins identified in uID–CD63 EV–treated cells are consistent with biologically relevant EV cargo.

## Discussion

Although affinity-based enrichments are widely used in proximity labeling and chemical biotinylation workflows, how enrichment efficiency depends on biotin stoichiometry and protein input remains poorly defined. This gap becomes increasingly consequential in advanced applications, including intercellular and *in vivo* labeling, which are often characterized by low protein input and limited biotinylation. In such settings, biotinylated targets represent only a minuscule fraction of the proteome, placing stringent demands on enrichment specificity and making it difficult to distinguish true signal from background binding. Robust assessment of enrichment performance under these conditions is therefore essential. To address this limitation, we establish a quantitative benchmarking framework that enables systematic evaluation of biotin enrichment performance under controlled conditions. By combining biotinylated *E. coli* protein standards with a human proteome background, this approach enables unambiguous discrimination of target and background proteins by MS. In this study, we defined enrichment efficiency using two metrics: (i) the number of target proteins recovered after enrichment and (ii) the fraction of the total protein signal represented by the recovered target proteins. We found that the number of recovered target proteins depends on both the level of biotinylation in the sample and the total protein input used for enrichment. In contrast, the fraction of recovered target proteins depended only on the level of biotinylation and was independent of the total protein input amount. Collectively, our results indicate that increasing the total protein input can be an effective strategy to improve the recovery of target proteins during biotin-based enrichment. We further indicate that, at least under the conditions tested, increasing protein input did not increase the ratio of target proteins recovered, as this proportion remained largely stable across input amounts. Specificity may instead be improved by raising the biotin stoichiometry of the sample, where feasible. In settings where a biotinylating enzyme is genetically introduced, increasing labeling duration or transduction efficiency may be beneficial to achieve higher biotinylation levels. Likewise, for chemical biotinylation, higher reagent concentrations or longer labeling times could improve the extent of labeling.

Our results suggest that biotin enrichments can reliably reflect relative differences in biotinylation between samples. At the same time, background proteins were stable between conditions with different biotin stoichiometries, indicating that biotin enrichments are a quantitative approach to trace differences of true analytes in differentially biotinylated samples. The fact that background proteins are not changing in abundance between samples with and without biotinylation highlights the utility of unlabeled control samples to distinguish between true analyte enrichment and non-specific background protein binding. Nevertheless, it should be noted that the samples in our experimental setup exhibit considerable differences in biotinylation levels. In the context of biotinylated EV uptake in recipient cells, differences between conditions are reduced, yet background protein levels remain stable. This finding indicates that false-positive identifications arising from non-specific binding are limited, even under conditions of low biotin content resulting from inefficient EV uptake.

We show that the quality of biotin enrichments is dependent on the affinity matrix used for the biotin enrichment. Here, we tested three different bead-based matrices which employed different biotin-binding proteins to enrich biotinylated proteins. In our hands, Pierce agarose beads coupled to neutravidin gave the best results, with more target proteins and a higher percentage of target proteins retrieved compared to Pierce magnetic and Resyn beads both coupled to streptavidin. This finding underscores that the choice of beads is critical but sometimes overlooked determinant of enrichment efficiency, a particularly important consideration when biotin stoichiometries of samples are low. However, it is also clear in the field that each bead type might have wide quality variation between different production lots, which influence the outcome of enrichments greatly ^25,26^. The lot variation of beads was an aspect we did not take into consideration in this work.

On-bead digestion is commonly used following protein-level enrichment for MS, since the high-affinity streptavidin-biotin interaction prevents efficient elution of target proteins. A drawback of this strategy is the partial proteolysis of streptavidin and other bead matrix components, which may contribute contaminating peptides to the dataset. The resulting highly abundant bead-derived peptides can dominate MS spectra and hamper detection of true analytes. Making beads resistant to protease digestion could therefore reduce this interference. We found that while the protease resistant beads do show a clear reduction in bead-derived peptide signal, the number of retrieved target analytes was slightly reduced compared to unmodified beads. We hypothesized that the harsh conditions applied to make the beads protease resistant could lead to a decrease in their binding capacity. This observation may be partly caused the use of data-independent acquisition (DIA). Because DIA gas-phase fractionates peptides for MS2 acquisition and many identifications rely solely on MS2 spectra, these measurements may be less susceptible to interference from a few highly abundant peptides. We therefore suggest that when dealing with a case of overloading of bead-derived peptides, considering DIA instead of data-dependent acquisition might be beneficial to alleviate the dynamic range problem of overloaded peptides. Although protease-resistant beads slightly reduced the number of recovered target proteins, they may still be advantageous when missing a few targets is not critical. Reduced proteolysis of biotin-binding proteins on beads does not only limits peptide overloading, potentially extending the lifespan of LC and MS systems, but also allows samples to be analyzed on more sensitive instruments with lower input requirements ideally enabling researchers to work with lower protein input samples.

We next assessed whether our findings on biotin enrichment efficiency extend to biotin-based labeling experiments conducted in a biologically relevant cellular setting. In the context of EV-mediated protein transfer, direct biochemical evidence for cargo delivery to recipient cells remains limited, hindering mechanistic insight into EV function. Using an EV transfer setup with distinct biotinylation levels in sender cells, isolated EVs, and receiver cells, we directly compared these samples to our biotinylated protein standards. We show that comparing biotinylated protein content of these 3 sample types to the biotinylated protein standards correlated well with their enrichment quality. While biotinylated sender cells and biotinylated EVs showed excellent enrichment with most proteins upregulated, receiver cells fed with biotinylated EVs compared to WT EVs showed upregulation of 14 proteins. The 14 proteins we identified are consistent with previous reports that EV uptake enhances the metabolic potential of receiver cells ^27,28^. Notably, these proteins rank among the highest-intensity biotinylated EV proteins after enrichment, highlighting that the method is operating near its detection limit in this setting. Achieving a more comprehensive characterization of EV-transferred proteins in recipient cells will require further improvements in the sensitivity of biotin enrichment workflows. Recent advances enabling enrichment from low protein inputs suggest a promising path toward increased detection depth and proteome coverage^29^. However, as experimental designs move toward progressively lower protein inputs and reduced biotinylation levels, distinguishing true biological signal from enrichment artefacts becomes increasingly challenging. In this context, our results underscore that a quantitative benchmarking framework is not merely supportive, but essential for rigorously assessing enrichment performance. By providing an accessible and scalable strategy to evaluate specificity under controlled and biologically relevant conditions, this framework offers a critical foundation for the reliable development and interpretation of emerging low-input biotinylation experiments.

## Conclusions

The expanding use of affinity-based enrichments in proximity labeling increasingly demands rigorous evaluation of enrichment performance, particularly for low-input or low-biotin biological samples such as intercellular or *in vivo* labeling experiments. We introduce a benchmarking strategy that quantitatively assesses enrichment efficiency under controlled conditions and demonstrates its applicability to biologically complex systems, including EV-mediated protein transfer. By providing a reliable metric to distinguish true enrichment from non-specific background, this framework enables systematic optimization of workflows and informs experimental design for challenging samples. As proximity labeling applications advance toward more demanding settings, this approach offers a practical and reproducible tool to guide method development and ensure interpretable results.

## Methods

### Mammalian cell culture

K562 were cultured in RPMI1640 with stable glutamine (Capricorn #RPMI-STA) supplemented with 100 U/ml Penicilin/streptomycin (Thermo Fisher # 15140122), 1x non-essential amino acids (Thermo Fisher # 11140050), 1 mM sodium pyruvate (Thermo Fisher # 11360070) and 10% (v/v) FBS. HeLa FlpINs were cultured in DMEM with stable glutamine and sodium pyruvate (Capricorn #DMEM-HPSTA) supplemented with 100 U/ml Penicilin/streptomycin (Thermo Fisher # 15140122) and 10% (v/v) FBS.

HeLa FlpIN cell stably expressing a TetR and uID-CD63 or uID-TSPAN3 were derived from the HeLa FlpIN cell line by transfection with pcDNA5/FRT/TO vector (Invitrogen) subcloned with Myc-uID-CD63 or Myc-uID-TSPAN3 and pOG44 (Invitrogen) and cultured in full media supplemented with 200 µg/mL hygromycin and 4 µg/mL blasticiding ^30^. CD63 or TSPAN3 localized uID activity of the resulting clones were validated after supplementation of biotin (10 min, 50 µM) using immunofluorescence (streptavidin-AF594 (1:5000, Invitrogen, S11227)), mouse-anti-Myc (1:500, 9E10, Santa cruz biotechnology, sc-40) followed by goat-anti-mouse-AF488 (1:1000, Invitrogen, A-11001).

### TurboID labelling

Cytosolic and GPI-anchored TurboID containing K562 cells were maintained in biotin-free RMPI media 4 days prior the experiment. Cells were labelled for 20h or 30 min with a final concentration of 100 uM Biotin at 37°C in a humidified incubator with 5% CO*2*. For efficient extracellular labeling, 1,5 mM ATP and 5 mM Vitamin B5 were added. After labelling, cells were washed 3x with PBS and lysed.

### Bacterial cell culture and lysis

DH5α Ecoli cells were thawed and incubated in LB (Broth) media overnight shaking at 37°C, 5% CO*2*. Grown cultures were harvested and washed three times with PBS.

### Lysis and protein extraction

The same lysis and protein extraction protocol was applied to K562 cells, *E. coli* cells, HeLa cells, HEP2G cells and isolated EV samples except otherwise state. All experimental steps were conducted on ice, except otherwise stated. Cell samples were pelleted at 400xg for 5 min and washed twice with PBS (Capricorn #PBS-1A). Cell samples were pelleted at 400xg for 5 min, flash frozen in liquid nitrogen and stored at -80°C. Frozen cell pellets or EV samples were then resuspended in of ice cold lysis lysis buffer(100 mM phosphate buffer, ph 7.1, 150 mM sodium chloride, 0.5 mM ethylenediaminetetraacetic acid (EDTA), 1 mM ethyleneglycol-bis(2-aminoethylether)-N,N,N’,N’-tetraacetic acid (EGTA), 1 mM magnesium chloride, 1% (v/v) nonyl phenoxypolyethoxylethanol (NP-40), 0.1% (w/v)sodium dodecyl sulfate (SDS), 0.4% (w/v) sodium deoxycholate (SDC)). Lysis buffer was complemented with cOmplete protease inhibitor cocktail (Roche #11836170001). Cell lysate was sonicated using a probe sonicator at 60% amplitude for 2x 1 min with 1 min break on ice in between. 70 µl of RNAseA and 70 µl of DNAse were added to the samples. Protein concentration was determined using bicinchoninic acid (BCA) assay (Thermo Fisher #A55864).

### Labeling of *E. coli* protein lysates

A total of 660 µL of *E. coli* protein lysate was biotinylated with Sulfo-NHS-SS-Biotin (Thermo Fisher Scientific) by adding 303.5 µL of a 10 mM Sulfo-NHS-SS-Biotin solution. The reaction was incubated for 2 hours at 4 °C and quenched by adding 1 M Tris-HCl (pH 7.5) to a final concentration of 50 mM. Samples were flash frozen in liquid nitrogen and stored at -80°C.

### Protease digestion resistant bead production

Protease digestion resistant beads were produced as previously described with slight adaptations ^31^. 500ul of neutravidin agarose beads were added to a 2ml falcon tube. Tube was centrifuged at 2500g for 2.5min and supernatant was discarded. Beads were washed with 1ml of PBS + Tween 0.1% (PBS-T). Tube was centrifuged at 2500g for 2.5min and supernatant was discarded. Beads were resuspended in 1.4ml of PBS-T at pH 13 containing 120mg of cyclohexanedione. Tube was incubated on a overhead rotator for 4h at room temperature. Tube was centrifuged at 2500g for 2.5min and supernatant was discarded. Tube was washed with 1ml of PBS-T. Tube was centrifuged at 2500g for 2.5min and supernatant was discarded. Beads were resuspended in 0.7ml of 4% formaldehyde in PBS-T. 0.7ml of 0.2M cyanoborohydride in PBS-T (solution is highly toxic). Tube was incubated on a overhead rotator for 2h at room temperature. Tube was centrifuged at 2500g for 2.5min and supernatant was discarded (check toxic chemical waist disposal procedures in your lab). Beads were washed with 1ml of 0.1M Tris-HCl at pH 7.5. Tube was centrifuged at 2500g for 2.5min and supernatant was discarded (according to toxic chemical waist disposal procedures in the lab). Beads were washed with 1ml of PBS-T. Tube was centrifuged at 2500g for 2.5min and supernatant was discarded. Beads were resuspended in a final volume of 0.5ml of PBS-T and kept in at 4°C.

### Agarose bead enrichment

For agarose bead enrichment, 20 µl of agarose beads (normal beads or protease resistant beads) were added to 1.5 ml Protein LoBind tubes and washed twice with 500 µl of lysis buffer, centrifuged at 2500xg for 2.5 min and supernatant was discarded. Protein lysates (500, 300, 100ug) for enrichment were added to the tubes and tubes were incubated for 3h at room temperature using an overhead rotator. After incubation samples were centrifuged at 2500xg for 2.5 min to collect beads at the bottom of the tube. Samples including beads were transferred to a microspin column (Biorad #732-6204). Beads were washed with the following buffers except otherwise stated. Beads were washed twice with 500 µl of each of the following buffers: 2% (w/v) SDS; 1% (w/v) SDC,1% (v/v) Triton X-100, 25 mM lithium chloride; 5 M sodium chloride; 50 mM ammonium bicarbonate (ambic); 30% (v/v) acetonitrile, 50 mM ambic. For supplementary figure 1a, washing conditions slightly differed. Harsh & organic condition refers to the standard wash condition just mentioned. Harsh condition refers to the standard wash condition without the 30% (v/v) acetonitrile wash. Control wash condition refers to three washes with 50 mM TEA, 150 mM NaCl, 5 mM EDTA, 0.1% SDS followed by 3 washes with 50 mM TEA, 150 mM NaCl, 5 mM EDTA. After washing, beads were transferred to a 0.5 ml protein LoBind Eppendorf tube by resuspending the beads on the filter in 200 µl of 50 mM ambic. Transferring step was repeated once to make sure all beads were transferred from filter to tube. Samples were centrifuged at 2500xg for 2.5 min and supernatant was discarded. Samples were washed twice with 500 µl of 50 mM ambic, centrifuged at 2500xg for 2.5 min and supernatant was discarded. Beads were resuspended in 40 µl of 50 mM ambic and 100 ng of trypsin (Promega # V5111) was added. On bead digestion was performed by digesting samples in a thermal shaker shaking at 1000 rpm and set at 37 ºC. After overnight incubation, another 100 ng of trypsin was spike into the samples and digestion was continued for another 3h. Digestion was stopped by adding formic acid to a final concentration of 2% (v/v). Tubes were centrifuged at 2500xg for 2.5 min and supernatant containing peptides was transferred to a new tube. Samples were frozen at -20 ºC until further processed by peptide cleanup.

### Pierce magnetic and resyn bead enrichment

For pierce magnetic and resyn bead enrichment, 20 µl of beads were added to 1.5 ml Protein LoBind tubes and washed twice with 500 µl of lysis buffer by placing them on a magnetic rack to separate beads from supernatant and supernatant was discarded. Protein lysates for enrichment were added to the tubes and tubes were incubated for 3h at room temperature using an overhead rotator. After incubation samples were placed on the magnetic rack and supernatant was discarded. Beads were washed twice with 500 µl of each of the following buffers: 2% (w/v) SDS; 1% (w/v) SDC,1% (v/v) Triton X-100, 25 mM lithium chloride; 5 M sodium chloride; 50 mM ammonium bicarbonate (ambic); 30% (v/v) acetonitrile, 50 mM ambic. Beads were resuspended in 40 µl of 50 mM ambic and 100 ng of trypsin (Promega # V5111) was added. On bead digestion was performed by digesting samples in a thermal shaker shaking at 1000 rpm and set at 37 ºC. After overnight incubation, another 100 ng of trypsin was spike into the samples and digestion was continued for another 3h. Digestion was stopped by adding formic acid to a final concentration of 2% (v/v). Tubes place on a magnetic rack and supernatant containing peptides was transferred to a new tube. Samples were frozen at -20 ºC until further processing by peptide cleanup.

### Immunoblotting

Approximately 20 µg of protein lysate was loaded per lane on a Criterion XT precast 1 mm, 12% Bis-Tris Acrylamide SDS-PAGE gel (Bio-Rad). The gel was initially run at 50 V for 5 minutes to ensure even sample distribution, followed by 2 hours at 100 V for protein separation. Proteins were then transferred to 0.2 µm PVDF blotting membrane (Transfer pack, Bio-Rad) using the Trans-Blot Turbo system (Bio-Rad) for 15 minutes at 1,3M and 25V. Membranes were stained with Ponceau S (Sigma) to verify protein transfer and washed with Tris-buffered saline with 1% Tween20 (TBS-T). For detection of biotinylated proteins, membranes were blocked with 3% BSA in TBST for 1–1.5 hours, followed by incubation with Streptavidin-HRP (1:2000, Cell Signaling Technologies) at 4 °C overnight with end-to-end rotation. The following day, membranes were washed three times with TBST and directly imaged using Pierce ™ ECL Plus Western Blotting Substrate (Thermo Fisher Scientific) on an Amersham Imager 600 (GE Healthcare).

### EV production and isolation

HeLa FlpIN WT, HeLa FlpIN UID-CD63 and HeLa FlpIN UID-TSPAN3 cells were seeded in three 15 cm dishes and grown till 70% confluency. Media was refreshed with 25 mL/dish of media depleted of EVs by ultracentrifugation (16 hours, 200.000 xg) and supplemented with 50 µM biotin. After 24 hours, the conditioned media was collected, cleared from dead cells (5 min, 2000 xg), and debri (15 min, 4000 xg).

The cells were washed three times with PBS and subsequently lysed using RIPA buffer complemented with cOmplete protease inhibitor cocktail (Roche #11836170001). Conditioned media was concentrated using tangential flow filtration (30 kDa, Vivoflow 50R, VF05H2) till 20 mL, and further concentrated using Amicon® Ultra-4 centrifugal filters (100 kDa, Merck Millipore Ltd. UFC910024) till 1000 µL. Next, EVs were isolated using size-exclusion chromatography using automated fraction collector (Izon) with 70 nm qEV columns. 2x 500 µL sample was run using the fractionation protocol: 2.4 mL buffer volume and 13 × 1mL fractions. The two first fractions were pooled and used for further analysis.

### EV characterization

To determine global size range of the pooled EV fraction, 10 µL was used for dynamic light scattering using the Prometheus Panta (NanoTemper). Protein content of EV fractions were analyzed using the micro BCA™ protein assay kit (Thermo Scientific, #23235)

### EV feeding

HepG2 cells were seeded on 12-well plates and grown to confluency. Growth media was replaced by media depleted of EVs (by ultracentrifugation, 16 hours, 200.000 xg) after which 30 µg of biotinylated EVs from HeLa FlpIN WT, HeLa FlpIN UID-CD63 or HeLa FlpIN UID-TSPAN3 were added and incubated for 16 hours. After the incubation period, media was removed, cells were washed 3 times with ice-cold PBS and lysed for downstream analysis.

### Biotin enrichment of EV related samples

Biotin enrichment of EV related samples was performed as agarose bead enrichments described above with differing protein input amounts. For sender cell enrichments, 500 µg of protein input was used. For isolated EV biotin enrichment, 50 µg of protein input was used. For receiver cells fed with biotinylated EVs, 45 µg of protein input was used.

### Whole EV lysate sample preparation

For whole EV lysate sample preparation sp3 protocol was followed with minor adaptations ^32^. In short, a volume of EVs corresponding to 5 µg protein input was lysed in lysis buffer (0.2% SDS, 0.5% SDC, 150 mM NaCl, 1% Triton-X. Lysis buffer was complemented with cOmplete protease inhibitor cocktail (Roche #11836170001). 4 µl of both SP3 bead types was added to each sample after washing 2 times with 500 µl of miliQ water. Sample volumes were supplemented with 100% Ethanol to reach a 80% Ethanol solution. Samples were incubated for 20min at room temperature with 1000rpm of shaking. Samples were then put on a magnet and supernatant was discarded. Samples were washed 2 times with 80% Ethanol in water, 2 times 100% acetonitrile, 2 time 50 mM ammonium bicarbonate in water. Beads were finally resuspended in 20 µl of 50 mM ammonium bicarbonate in water. Trypsin (Promega # V5111) was added in a trypsin to protein ratio of 1:25 (0.2 µg of trypsin per sample) and samples were on bead digested for 1h at 47ºC. After 1h of digestion, the same amount of trypsin was added and samples were digested for 1h at 47ºC. Supernatant was recovered to a fresh tube and beads were washed with 200 µl of 50 mM ammonium bicarbonate in water for a second recovery step. After second recovery was added to the fresh tube together with first recovery, samples were acidified to 2% formic acid concentration. Next, samples were subjected to peptide clean up.

### Peptide cleanup

Peptide samples were cleaned up using Oasis HLB (Waters # WAT058951) in combination with a vacuum manifold. In short, cartridges were activated by flushing them with 150 µl of acetonitrile twice. Cartridges were washed by flushing them with 150 µl of 2% (v/v) formic acid (FA). Samples were loaded onto cartridges and flushed through. Cartridges were washed with another 150 µl of 2% (v/v) FA. Peptides were eluted by flushing thorough 50 µl 50% acetonitrile in 2%(v/v) FA twice. Samples were dried using a speedvac and samples were resuspended in 2% FA for mass spectrometric analysis.

### Tandem mass spectrometric analysis

Generated peptides (entire sample for enrichment or approximately 1 µg otherwise) were analyzed using a Exploris 480 system (Thermo Fisher) mass spectrometer coupled to an Ultimate3000 UHPLC system (Thermo Fisher). Peptides were concentrated and desalted by using a PepMap™ Neo Trap Cartridge (5 mm × 0.3 mm, 5 μm, Thermo Fisher Scientific). Peptides were then separated by using an in house produced 50-cm analytical column packed with C18 beads (Poroshell 120 EC-C18, 1.9 μm; Agilent Technologies) with a 90 min gradient. Buffer A was 0.1% FA while buffer B was 0.1% FA in 80% acetonitrile. The gradient started at 4% buffer B, increased from 4% to 11% in 3 min, from 11% to 30% in 58 min, from 30% to 44% in 5 min, from 44% to 55% in 5 min, from 55% to 99% in 3 min, 99% wash-out for 5 min. Column was re-equilibrated to 4% buffer B in 10min before start of next run. The flow rate was set to 300 µl/ml and the column was heated to 50 ºC to reduce pressure. Peptides were ionized by using a spray voltage of 2kV and a capillary heated to 275 ºC. The mass spectrometer was operated in data-independent acquisition (DIA) mode. MS1 survey scans were performed over a mass range of 350-2000 m/z at a resolution of 120’000 with automatic gain control (AGC) target set to 100% or injection time of 54 ms. MS1 survey scans were followed by DIA in 46 variable-width isolation windows (m/z isolation windows are listed in Supplementary Table 1). Precursor selection was achieved quadropol. Precursor fragmentation was done using high-energy collisional dissociation (HCD) with stepped collision energy of 27%, 30%, 33%. MS2 spectra acquisition was done in a mass range of 200 to 2000 m/z at a resolution of 30’000 at am AGC target of 3000% and a maximum injection time of 54 ms.

### Database searching

All data was searched by using DIA-NN 2.0 in library free mode ^33^. For human-*E*.*coli* mixed samples reviewed human and *E*.*coli* protein fasta files were obtained from uniport.org and merged into one fasta file. For human alone EV related samples reviewed human protein fasta files from uniport.org were used. Predicted spectral libraries were generated in DIA-NN using the beforementioned fasta files in a separate step. Raw files were then searched with the generated spectral libraries with precursor generation turned off. Normalization was disabled Screenshots of DIA-NN settings are added in supplementary information (Supplementary Figure 7). Data was filtered for at least 2 precursors per peptide. Protein intensities were aggregated from precursor intensities using the iq package implemented in the protti package in R ^34–36^.

### Data analysis

Data analysis and data visualization was performed in R ^37^. Imputation was performed using the protti package with minimal imputation, applying a standard distribution of values around the 5th percentile of protein intensities ^35^. Differential abundance calculations were performed with limma package implemented in the protti package in R ^35,36,38^.

## Acknowledgments

This work received funding support from NWO (ENW XL project OCENW.M.22.233) and the NWO-funded Netherlands Proteomics Center through the National Road Map for Large-scale Infrastructures program XOmics (Project 184.034.019). Additionally, this work was also supported by CROSSTALK (grant agreement no. 101075975).

## Data availability

Raw data of proteomics files are publicly available on PRIDE with the accession number PXD077873^39^. All the R scripts used in this study and any additional information required to reanalyze the data reported in this work are available upon request to the Lead contact.

## Contributions

Conceptualization: C.Z., M.C.T.S., J.B., F.J.V., M.A., K.S.; Data curation: C.Z., M.C.T.S., J.B; Formal analysis: C.Z., M.C.T.S., J.B; Funding acquisition : F.J.V., M.A., K.S.; Investigation: C.Z., M.C.T.S., J.B.; Methodology: C.Z., M.C.T.S., J.B., T.W.; Supervision: F.J.V., M.A., K.S.; Visualization: C.Z., M.C.T.S., J.B; Writing-original draft: C.Z., M.C.T.S., J.B., K.S.; and Writing - review & editing: C.Z., M.C.T.S., J.B., F.J.V., M.A., K.S.

## Supplementary Information

The supplementary information file contains additional Figures S1-S7 and table S1, as referenced in the text. Supplementary tables 2-4 are provided and include:

- protein intensities of biotin enrichments of 500ug of biotin standard samples shown in Figure 2 (Supplementary table S2)
- protein intensities of biotin enrichment with decreasing input amounts shown in Figure 3 (Supplementary table S3)
- differential abundance calculation of sender cell, EV and receiver cell enrichments shown in Figure 4 (Supplementary table S4)

## Notes

### Competing Interest Statement

The authors have declared no competing interest.

