## Supplementary Figures 1-7 and Supplementray Table 1 for "Stoichiometry-dependent specificity in biotin enrichment: a benchmarking framework for proximity labeling proteomics"

Netherlands.

^b^Netherlands Proteomics Center, Utrecht, The Netherlands.

^c^Department of Cell Biology, Neurobiology and Biophysics, Utrecht University, Padualaan 8, 3584CH Utrecht,, The Netherlands.

^#^These authors contributed equally to this work.

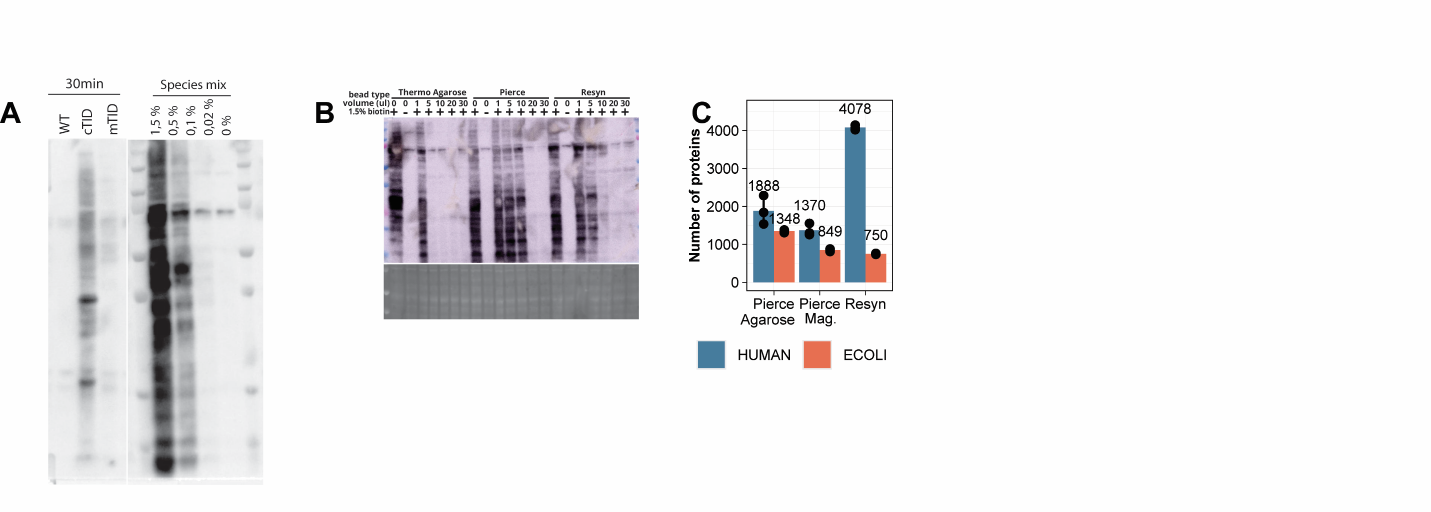

Supplementary Figure 1. Effects of wash conditions and bead types on enrichment efficiency.

**A)** Western blot of cell lysates of cytosolic (cTID) or membrane-localized TurboID (mTID) samples after 30 min labeling, alongside biotinylated benchmark standards. Each lane contains 20 µg of the respective sample. Biotinylated proteins were detected with streptavidin–HRP. **B)** Top: Westernblot of 500ug of 1.5% biotinylated *E. coli* lysate in human background was depleted with different volumes of either Pierce agarose, Pierce magnetic or Resyn beads. 20ug of initial sample was loaded on the WB. Biotinylated proteins are detected using streptavidin-HRP. Bottom: Ponceau S staining shows total protein loading of the corresponding blot. **C)** Bar plot showing the number of *E. coli* (red, target) and human (blue, background) proteins detected by LC–MS after enrichment of 500 µg of 1.5% biotinylated *E. coli* lysate in human background using different bead types.

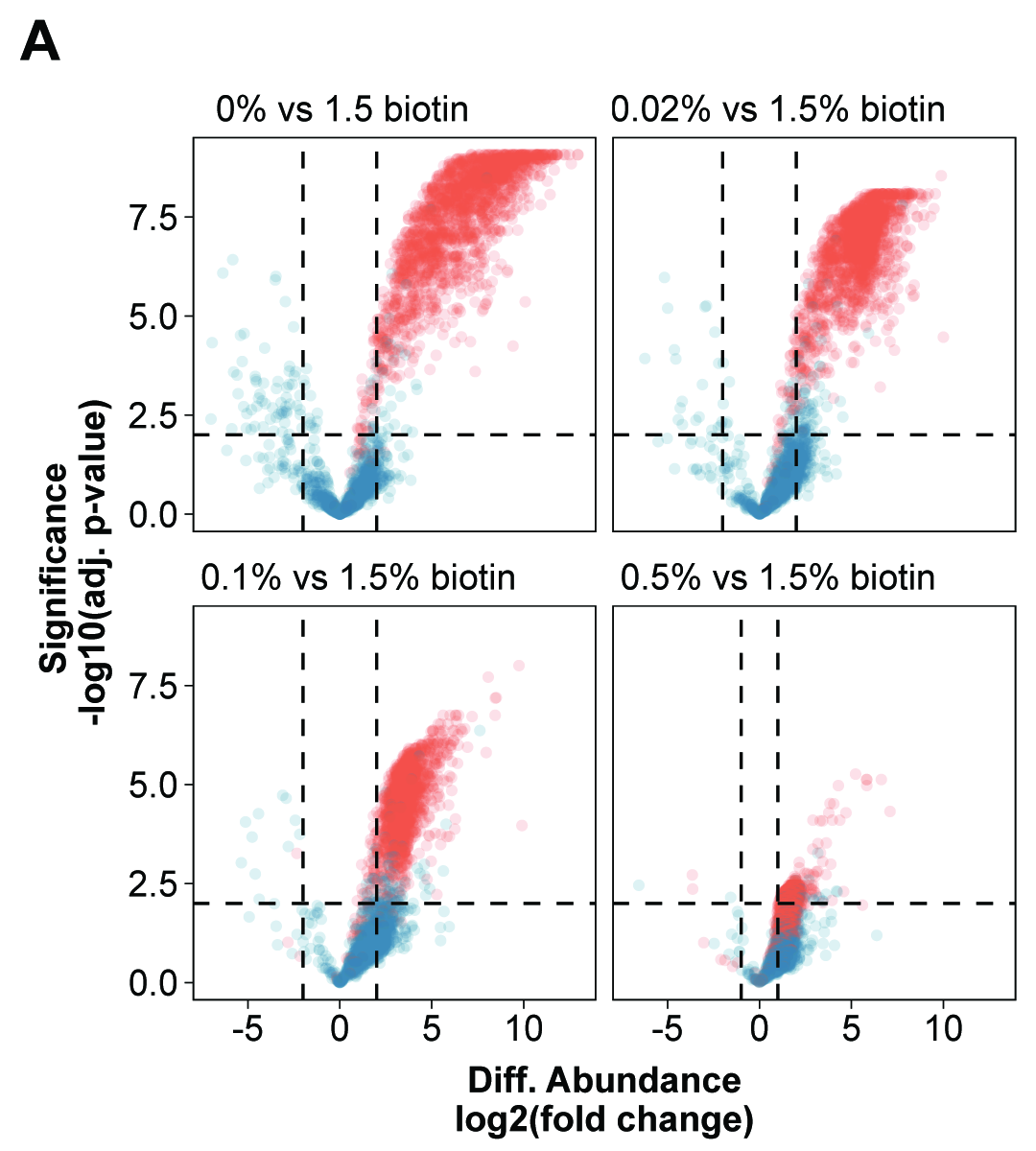

Supplementary Figure 2. Change of human background proteins across different biotinylation percentages.

**A)** 500ug of input sample at different biotinylation percentages was enriched with Pierce agarose beads (data shown previously shown in Figure 1 B and E). Volcano plots showing differential protein abundance versus –log10(adjusted p-value) for pairwise comparisons of biotinylation percentages of *E. coli* (red, target) and human (blue, background) proteins. P-values were calculated using a moderated t-test from the limma package, and adjustment for multiple testing was performed using the Benjamini-Hochberg method. Horizontal lines indicate an adjusted p-value threshold of 0.01. Vertical lines indicate log2 fold change thresholds: 2 for comparisons of 0, 0.02, and 0.1%, and 1 for the 0.5% comparison.

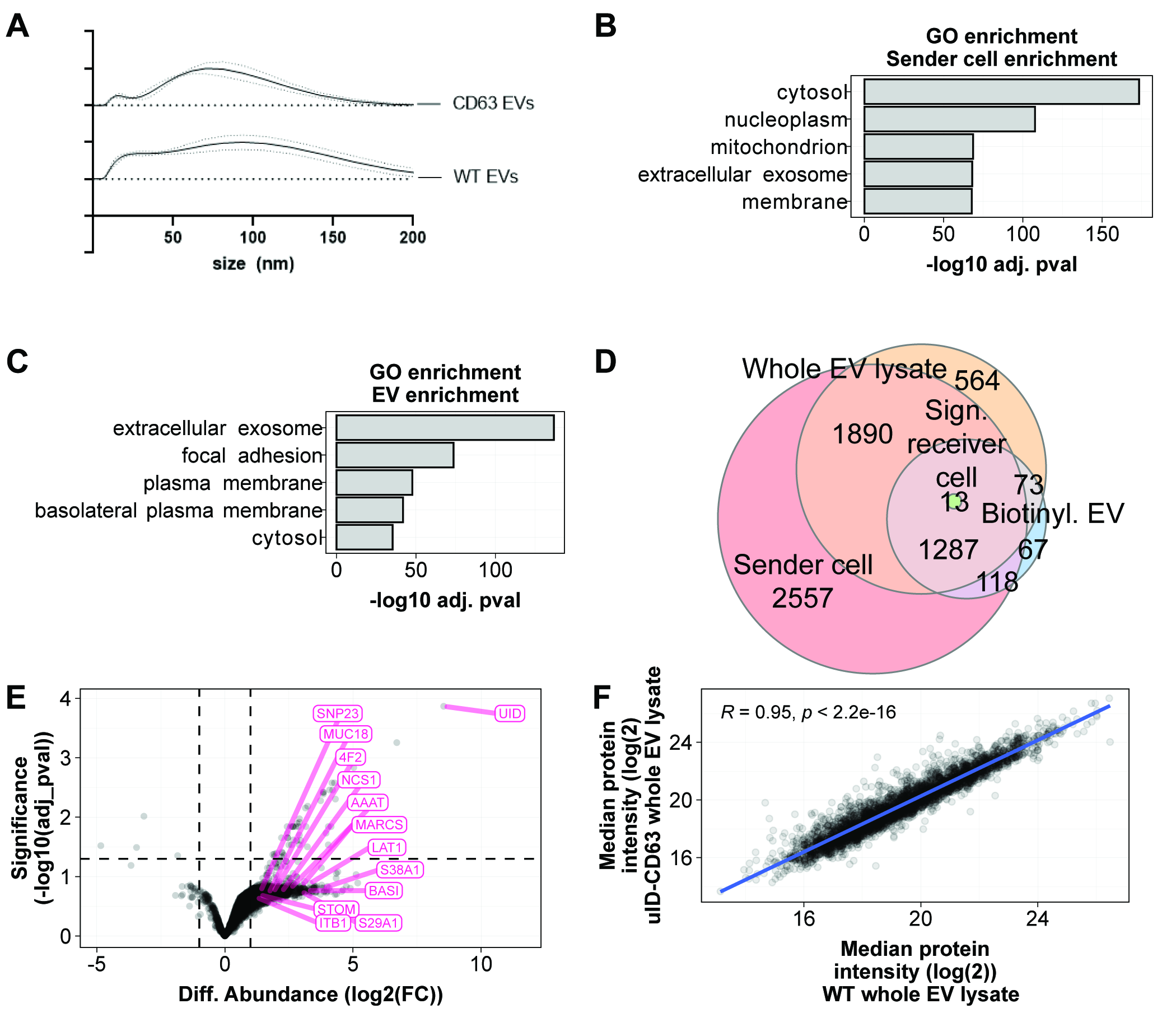

Supplementary Figure 3. Biotinylated sender cells and EVs are specifically enriched for EV proteins.

**A)** Particle size distribution profiles of CD63 EVs and wild-type (WT) EVs measured by dynamic light scattering. The x-axis shows particle diameter (nm) and the y-axis shows relative particle number (arbitrary units). **B&C)** Gene Ontology (GO) enrichment analysis of proteins significantly upregulated in uID-CD63 compared to wild-type (WT) samples (see volcano plots in Figure 4C). Significantly upregulated proteins were tested against the whole human proteome as background. Enrichment was assessed using Fisher’s exact test. Enriched GO terms in biotin-enriched sender cells (B) and in biotin-enriched EVs (C). **D)**Venn diagram showing overlap of proteins detected in sender cell enrichment (red circle), biotinylated proteins detected in EVs (light blue circle), biotinylated proteins found to be upregulated in receiver cells after feeding with biotinylated EVs (green circle) and proteins found in whole EV lysates (orange circle). **E)** Volcano plot showing differential protein abundance versus significance (–log10(adjusted p-value)) of whole EV lysate (D). Comparisons are always CD63-UID versus WT. Proteins significantly upregulated in receiver cell enrichment are highlighted in pink. **F)** Scatter plot showing the correlation of median protein intensities between wild-type (WT; x-axis) and uID-CD63 (y-axis) samples derived from whole EV lysates. Each point represents a quantified protein. The Pearson correlation coefficient is displayed at the top of the plot, indicating the strength of the linear relationship between the two conditions.

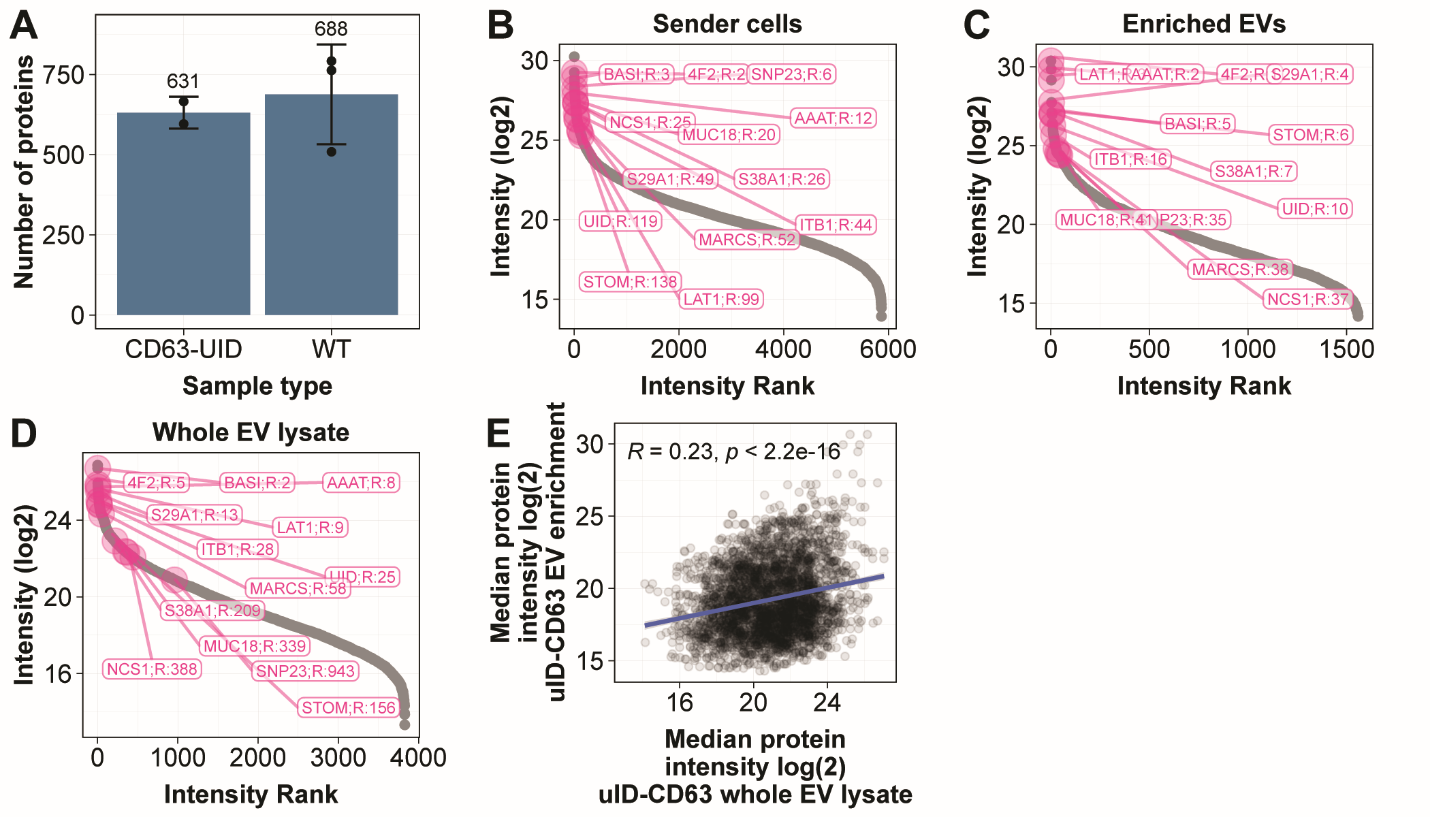

Supplementary Figure 4. Proteins enriched in receiver cells treated with biotinylated EVs correspond to high-ranking proteins in EV enrichment.

**A)** Number of proteins retrieved after biotin enrichment of receiver cells fed with EVs biotinylated by CD63-UID construct or with WT EVs. **B-D)** Rank plots of sender cells and EVs, showing log2 protein intensity (y-axis) versus intensity rank (x-axis). Panels B&C: biotin-enriched samples of sender cell lysates (B) and isolated EVs (C); panel D: whole EV lysates. Proteins significantly upregulated in receiver cells after biotinylated EV treatment are highlighted in pink. **E)** Scatter plot showing the correlation of median protein intensities between samples derived from uID-CD63 whole EV lysates (x-axis) and samples derived from biotin enriched uID-CD63 EV (y-axis). Each point represents a quantified protein. The Pearson correlation coefficient is displayed at the top of the plot, indicating the strength of the linear relationship between the two conditions.

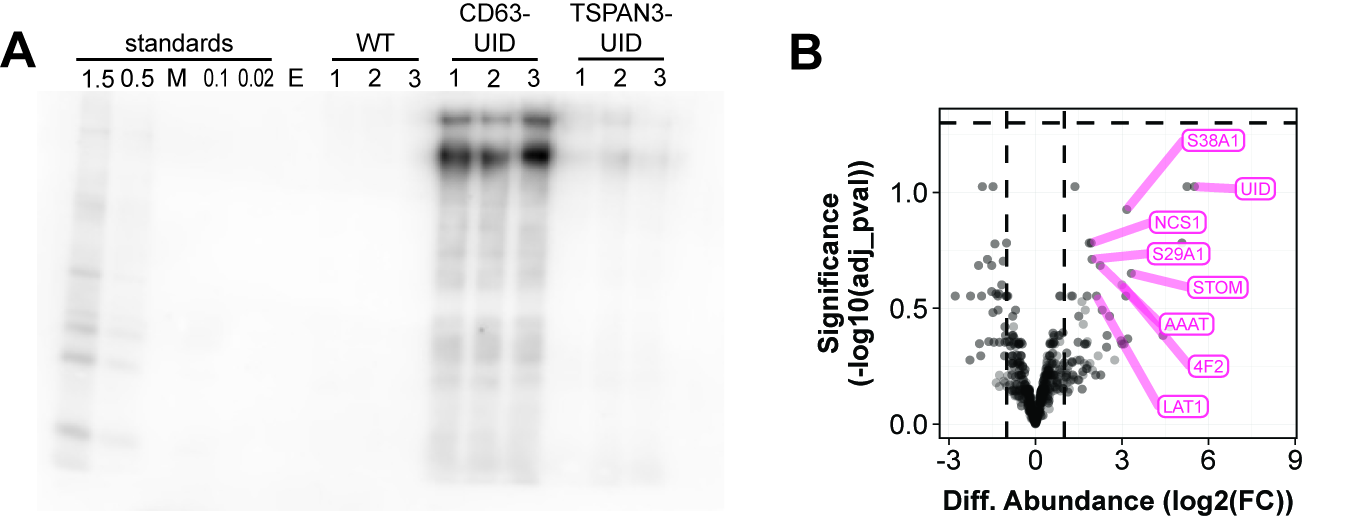

Supplementary Figure 5. Decreased biotin content of UID-TSPAN3 EVs leads to no detectable enrichment of biotinylated proteins after feeding to receiver cells.

**A)** Western blot of isolated EVs alongside biotinylated benchmark standards. Each lane contains 20 µg of the respective sample. Biotinylated proteins were detected with streptavidin–HRP. WT referres to EVs isolated from WT HeLa cells, CD63-UID refers to EVs isolated from HeLa cells genetically modified with CD63-UID construct, TSPAN3-UID refers to EVs isolated from HeLa cells genetically modified with TSPAN3-UID constructs. Letter E marks empty lanes. **B)** Volcano plot showing differential protein abundance versus significance (–log10(adjusted p-value)) of receiver cells fed with TSPAN3-UID containing EVs. Comparisons is TSPAN3-UID versus WT. Proteins significantly upregulated in receiver cell enrichment fed with CD63-UID containing EVs are are highlighted in pink.

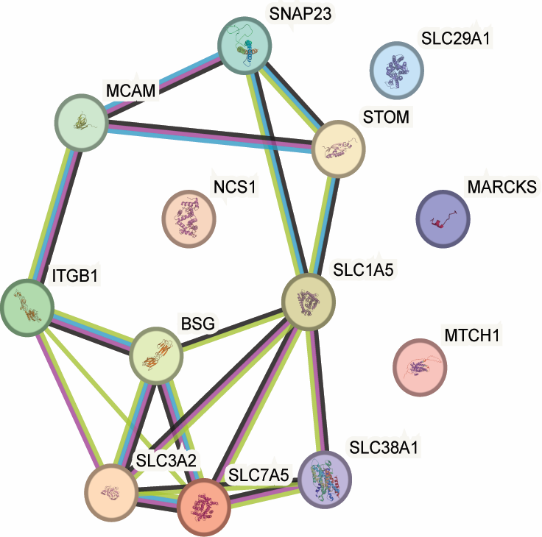

Supplementary Figure 6. Proteins significantly upregulated are closely related.

STRING network analysis of proteins significantly upregulated in receiver cells following biotinylated EV treatment. Only interactions with a confidence score ≥ 0.4 are shown.

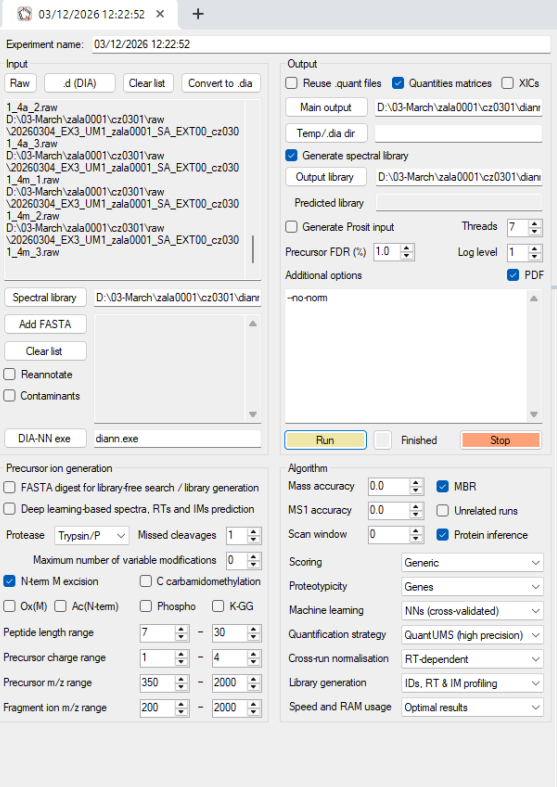

Supplementary Figure 7. DIA-NN settings used for searches.

Supplementary table 1. DIA window sizes used for DIA acquisition.

m/z range

399-407

406-414

413-420

419-426

425-432

431-438

437-444

443-450

449-455

454-461

460-467

466-472

471-479

478-484

483-490

489-496

495-502

501-509

508-516

515-522

521-529

528-536

535-543

542-550

549-558

557-565

564-574

573-583

582-592

591-602

601-613

612-624

623-637

636-649

648-663

662-677

676-694

693-712

711-732

731-756

755-782

781-813

812-851

850-906

905-1000

999-1650
